# Single molecule fluorescence *in situ* hybridisation for quantitating post-transcriptional regulation in *Drosophila* brains

**DOI:** 10.1101/128785

**Authors:** Lu Yang, Joshua S. Titlow, Darragh Ennis, Carlas Smith, Jessica Mitchell, Florence L. Young, Scott Waddell, David Ish-Horowicz, Ilan Davis

## Abstract

RNA *in situ* hybridization can be a powerful method to investigate post-transcriptional regulation, but analysis of intracellular mRNA distributions in thick, complex tissues like the brain poses significant challenges. Here, we describe the application of single-molecule fluorescent *in situ* hybridization (smFISH) to quantitate primary transcription and post-transcriptional regulation in whole-mount *Drosophila* larval and adult brains. Combining immunofluorescence and smFISH probes for different regions of a single gene, i.e., exons, 3’UTR, and introns, we show examples of a gene that is regulated post-transcriptionally and one that is regulated at the level of transcription. We also show that the method can be used to co-visualise a variety of different transcripts and proteins in neuronal stems cells as well as deep brain structures such as mushroom body neuropils. Finally, we introduce the use of smFISH as asensitivealternative to conventional antibody labelling to mark specific neural stem cell populations in the brain.

## 1. Introduction

The central nervous system (CNS) consists of an extraordinary number and diversity of cells, most of which are derived from a relatively small number of neural stem cells. Biochemical methods have been instrumental in elucidating post-transcriptional regulatory mechanisms, but these methods typically involve dissociation and homogenization of tissues [1] and therefore offer only limited spatial information. In this paper, we describe an RNA *in situ* hybridization (ISH) method that can provide effective measurements of gene expression within the spatial context of a whole *Drosophila* brain.

Single molecule fluorescence *in situ* hybridization (sm-FISH) has revolutionized the potential of RNA ISH by enhancing sensitivity and probe penetration [2-3]. The state-of-the-art smFISH technique uses 25-48 individual fluorescently labeled DNA oligonucleotide (oligo) probes approximately 20 bases long, tiling a region of a target transcript. The use of short oligos improves probe penetration while the relatively large number of probes allows single molecules to be detected as bright foci, which are easily distinguishable from background fluorescence generated by nonspecific labeling [4-5]. The use of directly-coupled fluorochromes to the oligos eliminates the signal amplification step in the classical DIG-Tyramide-FISH protocol. So far the smFISH method has enabled the study of gene regulation in single-cell organisms, *in vitro* cell culture systems [6-7], and in *Drosophila* oocytes, embryos, and the larval neuromuscular junction [8-11]. However, smFISH is still dependent on the development of specific conditions for individual tissue types, and the use of smFISH on thick tissue such as the larval or adult brain has remained particularly challenging.

Here, we have adapted the smFISH method to whole-mount *Drosophila* brain tissues, providing quantitative information at sub-cellular resolution. We demonstrate how simultaneously labeling the intron and exon of a gene with separate smFISH probes can be used to quantitate primary transcription in comparison to post-transcriptional cytoplasmic levels of mRNA (section 4.2). Combining smFISH with antibody labeling of the protein encoded by the same gene provides a direct measure of post-transcriptional regulation (section 4.4). Finally, we also show that smFISH can be used as a marker to identify specific cell types (section 4.5).

## 2. Materials and reagents

### 2.1 Probe design and preparation

The minimum number of probes that generated an acceptable signal-to-noise ratio in the larval and adult brain tissue is 30 for the genes presented. However, this number greatly depends on the native expression level of the specific transcript, binding affinity of the probes, and the type of dye. Several dyes are available for labeling smFISH probes. In our hands, Quasar-570, Quasar-670, and Atto-647N provide an effective signal to noise ratio in the *Drosophila* brain, whereas fluorescein does not. Multiple strategies exist for synthesizing probes conjugated to a large selection of different dyes [12-14]. Here StellarisTM smFISH probes were purchased from LGC BioSearch Technologies (California, USA). A set of oligonucleotide probes specific to the gene of interest was created using the web-based probe designer https://www.biosearchtech.com/stellaris-designer.

If no single region of the gene is sufficient in length, probes can be generated from multiple combined regions of the same gene. This can be particularly useful when designing a probe set against intronic regions of the gene of interest. We recommend downloading the probe sequence and assessing the probe sequence specificity (e.g., using the free, web-based BLAST program https://blast.ncbi.nlm.nih.gov). A negative control is also essential for distinguishing smFISH signal from background noise and non-specific binding. We find that the best negative controls are those of a biological nature. Here we demonstrate the use of smFISH probes targeting YFP in a wild-type background as a negative control (Figure 2G-G”’). For endogenous genes, the smFISH probes could be tested in a transcript-null mutant or an RNAi knockdown for the gene of interest.

### 2.2 Reagents and buffers for smFISH

- 0.3% PBSTX (1x phosphate buffered saline (PBS) with 0.3% TritonX)
- 0.3% PBST (1x PBS with 0.3% Tween-20)
- Fixation buffer (4% formaldehyde in 0.3% PBSTX)
- WASH buffer (10% v/v deionised formamide^1^ in 2x saline sodium citrate (SSC) solution^2^)
- Hybridization buffer^3^ (10% v/v deionized formamide, 10% v/v of 50% Dextran Sulphate solution (Millipore) (final dextran concentration - 5%) in 2x saline sodium citrate (SSC) solution

### 2.3 Reagents and buffers for immunofluorescence with smFISH

- Blocking buffer (0.1% goat serum, 1% glycine in 0.3% PBSTX^4^)
- Primary antibodies, mouse anti-Dlg1 (Developmental Studies Hybridoma Bank #4F3, 1:500), rat anti-Dpn (Abcam ab195173, 1:500), rat anti-mir (J. Knoblich lab, 1:100), guinea pig anti-Ase (J. Knoblich lab, 1:50), goat anti-HRP conjugated to Dylight405 (Jackson ImmunoResearch, 1:100)
- Bovine serum albumin (nuclease-free)

### 2.4 Drosophila strains

Fly stocks were maintained at 25°C on 12hr light:dark cycle. The following genotypes were used: Wild type-Canton S, Dlg1::GFP, UAS-mcd8:GFP (Bloomington Drosophila Stock Center no. 5137)[15], Pros-Gal4, Imp::GFP (Bloomington Drosophila Stock Center no. 41500)[16].

## 3. Protocols

### 3.1 smFISH protocol

1. An overview of the smFISH workflow is illustrated in Fig. 1A. Dissect brains from 3rd instar larvae or adult flies in Schneider’s medium. To minimize tissue damage in larva dissections, we recommend a scissors dissection method as opposed to only using forceps (Fig. 1B). Adult brains can be dissected using standard techniques for fixed or live brain imaging [17]. After dissection, all steps in the smFISH procedure are identical for larval and adult brains.
2. Fix brains in fixation buffer for 20 min.
3. Quickly rinse brains 3 times with 0.3% PBST.
4. Wash brains 3 times for 15 min each at 25°C with 0.3% PBST^5^.
5. Incubate brains in wash buffer for 5 min at 37°C.
6. Incubate brains in hybridization buffer with the appropriate probe concentration^6^ at 37°C for 8-15 hours with gentle rocking^7^. Samples should be protected from light for all subsequent steps (see Fig. 1A for a setup of light-proof sample chamber). Antibodies can be included in this step at concentrations typically used for immunofluorescence (See Section 2.3 for specific concentrations).
7. Rinse sample 3 times in wash buffer.^8^
8. Wash brains 3 times for 15 min each time with wash buffer at 25°C. The nuclear stain DAPI (2 μg/ml) can be included during the penultimate wash. Secondary antibodies should also be included in this step for experiments involving immunofluorescence, and the wash should be extended to 45mins.
9. Wash sample for 10 min at 25°C with 0.3% PBST. This step prevents the brains from adhering to the inside wall of the pipet tips during mounting.
10. Proceed to sample mounting (section 3.2).

**Figure 1.**
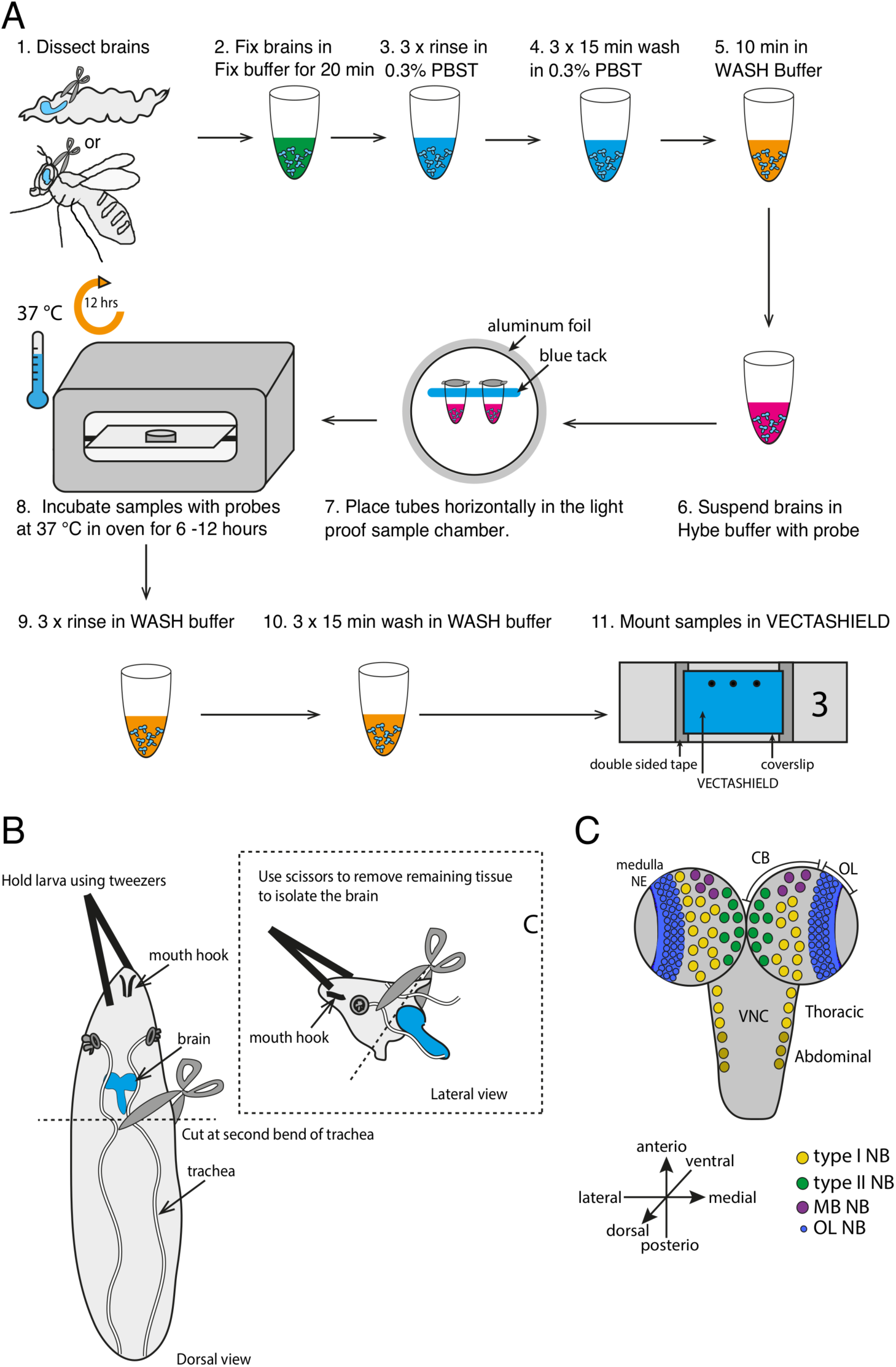
Overview of smFISH, larval brain dissection, and orientation. A Overview of the smFISH protocol for brain tissue. After brains are dissected from third instar larvae or adult flies, samples are fixed in 4% formaldehyde. After series of washes, brains are incubated in hybridization buffer containing the probe mixture targeting the gene of interest. Hybridization step takes place at 37°C for between 10-15 hours in a light protected environment. Following hybridization, samples undergo a further three washes and can then be mounted for imaging with the anti-fade mounting medium. B Scissor dissection method for larval brains. Immobilize larva by gently holding the tip of the larval head using a pair of dull tweezers. Then cut the larva in half at approximately the position of the second bend of the trachea tubes. The brain should be exposed from the remaining tissue and can be isolated using scissors. This dissection method reduces tissue damage in comparison to isolating the brain by pulling the larvae apart using two pairs of tweezers. C Distribution of different types of neuroblasts in the larval brain. Brain should be mounted in the orientation that is suitable for the purpose of the experiment.

### 3.2 Sample mounting

After the last wash, brains are transferred to a dissection dish by pipetting, and any unwanted tissues can be removed at this stage (it may be easier to remove some of the more closely attached connective tissues and imaginal discs from the brain at this stage rather than during the initial dissection because fixation in formaldehyde hardens the tissues and makes it easier to remove without damaging the samples). Place a coverslip on top of the mounting stage - a mounting stage can be made by simply taping two slides together with double sided tape.

1. Transfer brains to the coverslip using a 100μl pipet. Remove excess liquid from coverslip.
2. Pipet 20μl Vectashield^®^ (Vector Laboratories Ltd.) anti-fade mounting medium onto the coverslip to sufficiently cover the brains.
3. Align brains in a straight line while taking care to orient the tissue to be imaged as close to the coverslip as possible.^9^
4. Gently lower the pre-prepared micro slide^10^ to the coverslip, making sure the samples are positioned near the midline between the double-sided tape on each side.
5. Seal the coverslip with nail varnish. Care should be taken to store the slides in the dark at -20°C, as signal to background decreases over time.
6. Image slide with scanning confocal (higher quality) or spinning disk confocal (higher throughput) microscope. Images used in the current manuscript were acquired using an Olympus Fluoview FV1000 microscope with 40x 1.3 NA Oil UPlanFLN and 60x 1.35 NA Oil UPlanSApo objectives (Fig. 2B-B”’and Fig. 3-6), Zeiss LSM-880 with 60x 1.4NA Oil (Fig. 4D-D”’), or Perkin Elmer UltraView Spinning Disk with 60x 1.35 NA Oil UPlanSApo objectives (Fig. 4F-G”’).

**Figure 2.**
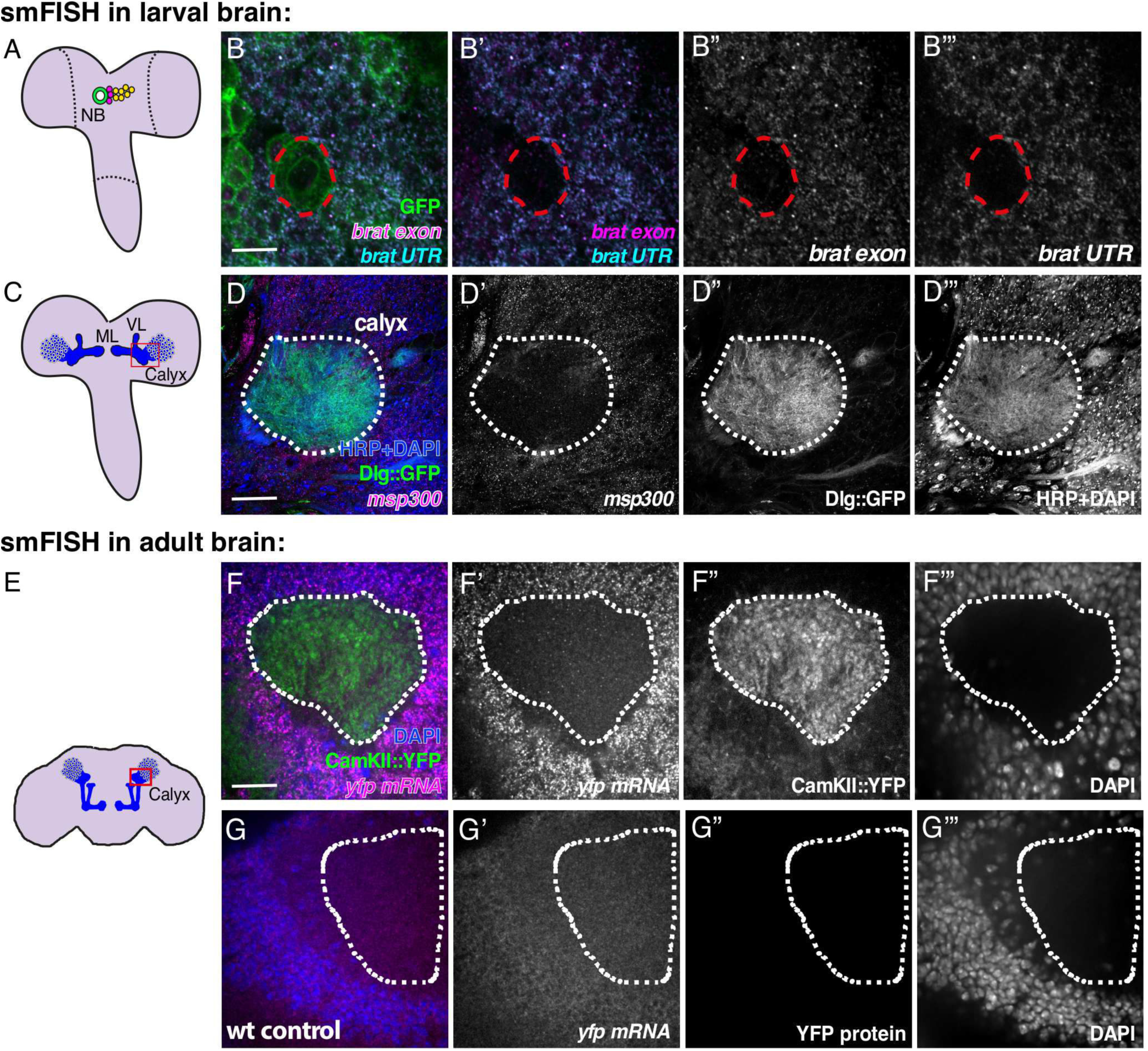
Simultaneous detection of multiple RNA species using smFISH. A Schematic of the 3rd instar *Drosophila* larva brain illustrating the position of a neuroblast lineage. B-B”’ Exemplar image of smFISH in the third instar larval brain generated using two sets of probes; one targets the exon region of the transcripts encoding RNA binding protein *brain tumor (brat)* (magenta) and the other targeting the 3’UTR of the same gene (cyan). Neuroblasts and progeny are labeled by driving the expression of membrane-tethered GFP using pros-gal4. C Schematic of the 3rd instar *Drosophila* brain illustrating the position of mushroom body lobes, calyx, and Kenyon cell nuclei. D-D”’ Detection of *msp-300* mRNA in the calyx of larval mushroom bodies. The mushroom body is identified by the expression of Dlg1 protein coupled to GFP, the axon bundle, and cell nuclei are labeled with HRP and DAPI, respectively. Scale bars represent 50μm. E Schematic of the adult *Drosophila* brain illustrating the position of mushroom body lobes, calyx, and Kenyon cell nuclei. F-F”’ Detection of *CaMKII::YFP* mRNA with a smFISH probe targeting the *YFP* mRNA sequence, and CaMKII::YFP protein in the adult mushroom body calyx (dotted line). G-G”’ Negative control showing the YFP smFISH probe in a wild-type adult brain. Abbreviations: NB-neuroblast, ML-medial lobe, VL-ventral lobe, wt-wild type.

**Figure 3.**
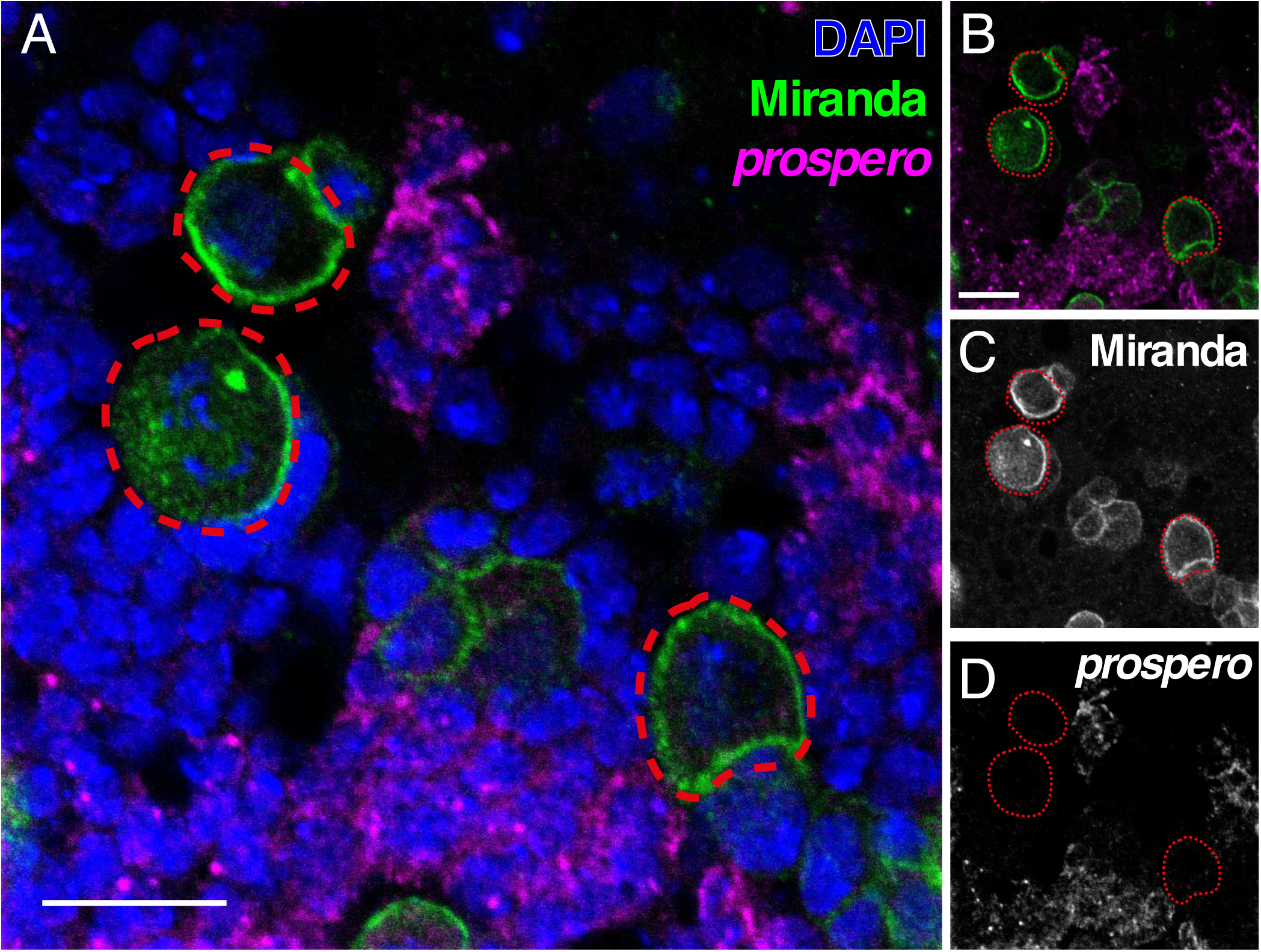
Simultaneous detection of RNA and protein using smFISH in conjunction with antibody staining A The smFISH protocol is compatible with conventional antibody staining. An exemplary image showing simultaneous detection of *Prospero* RNA and Miranda protein. B-D Image without nuclear DAPI stain Prospero RNA and Miranda protein (B) and the respective greyscale images (C-D). Scale bar represents 10μm (A) and 5μm (B-D).

**Figure 4.**
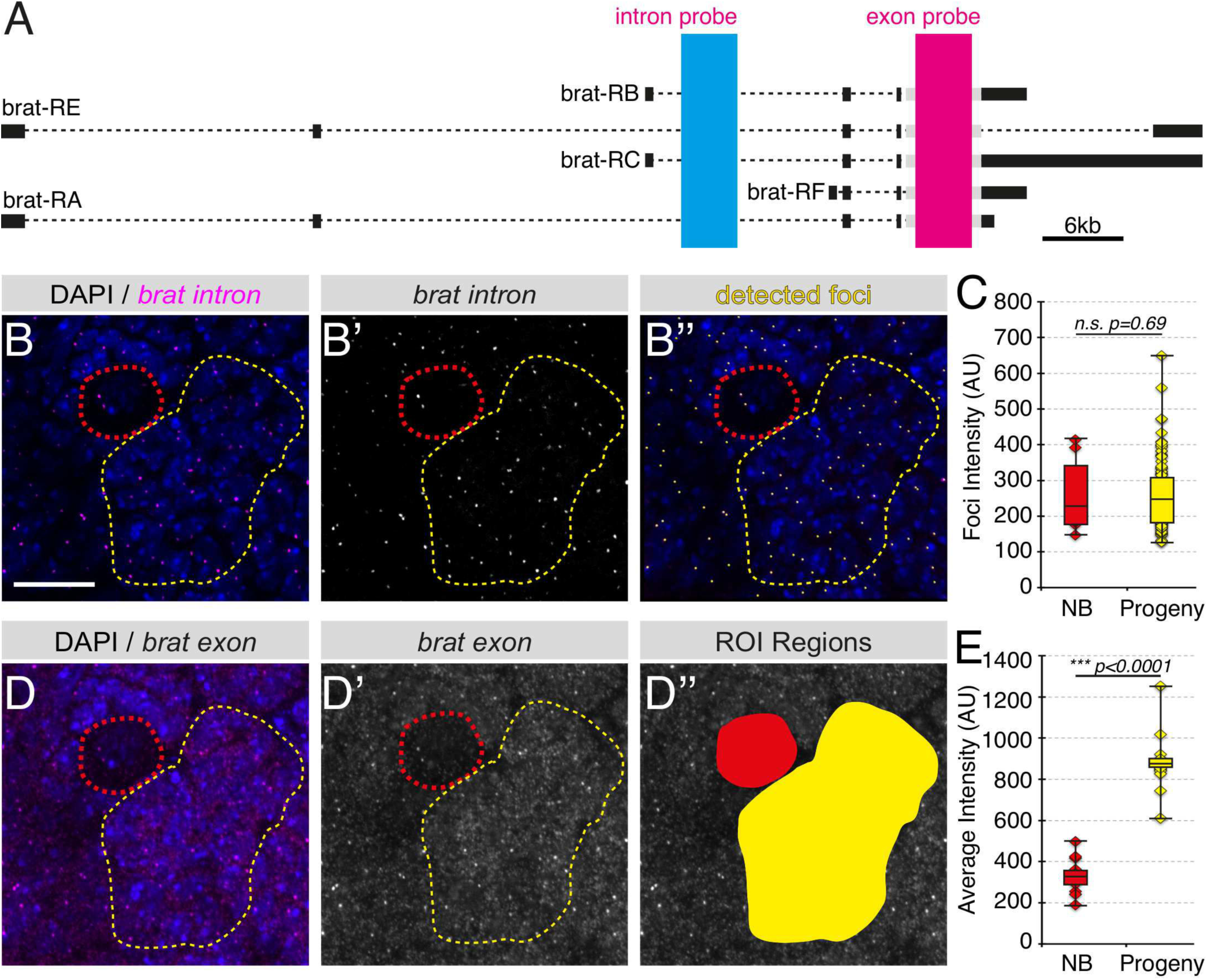
Distinguishing transcriptional versus post-transcriptional regulation with smFISH. A Location of the probe targeting the exon (magenta) and intron (blue) region of *brat* transcripts. B-B” Quantitative analysis of *brat* transcription level in neuroblast (red outline) and neuroblast progeny (yellow outline). The smFISH signal showing primary transcription foci in B and B’and automatic detection of foci using the “Spots” tool in the Imaris image analysis software (B”). C Average intensity of transcription foci in neuroblast and progeny is not significantly different, suggesting *brat* RNA is transcribed at approximately the same level in both cell types (neuroblast: n= 6 neuroblasts/brain, 3 brains; neuroblast progeny: n=100 foci, 3 brains). D-D” Quantitative analysis of total *brat* RNA in neuroblast and neuroblast progeny. The smFISH signals of textitbrat RNA detected by the exon probe is shown in D-D’and the region of interest selected for average intensity analysis is shown in D". Statistical analysis shows the level of total textitbrat transcripts is significantly increased in neuroblast progeny (neuroblast: 6 neuroblast/brain, 3 brains; progeny: 6 clusters/brain, 3 brains). Scale bar represents 10μm.

### 3.3 Statistical analysis

Datasets for average signal intensity or number of foci were tested for normality using the Shapiro-Wilk normality test. Data that deviated significantly from normal (p<0.05; Figs. 4C and 4E) were compared using the Wilcox rank sum test. Data with normal distributions (Figs. 5C and 5E) were compared using one-way ANOVA with Tukey post-hoc test. All statistical analyses were performed with R (version 3.3.2 in Jupyter Notebook).

**Figure 5.**
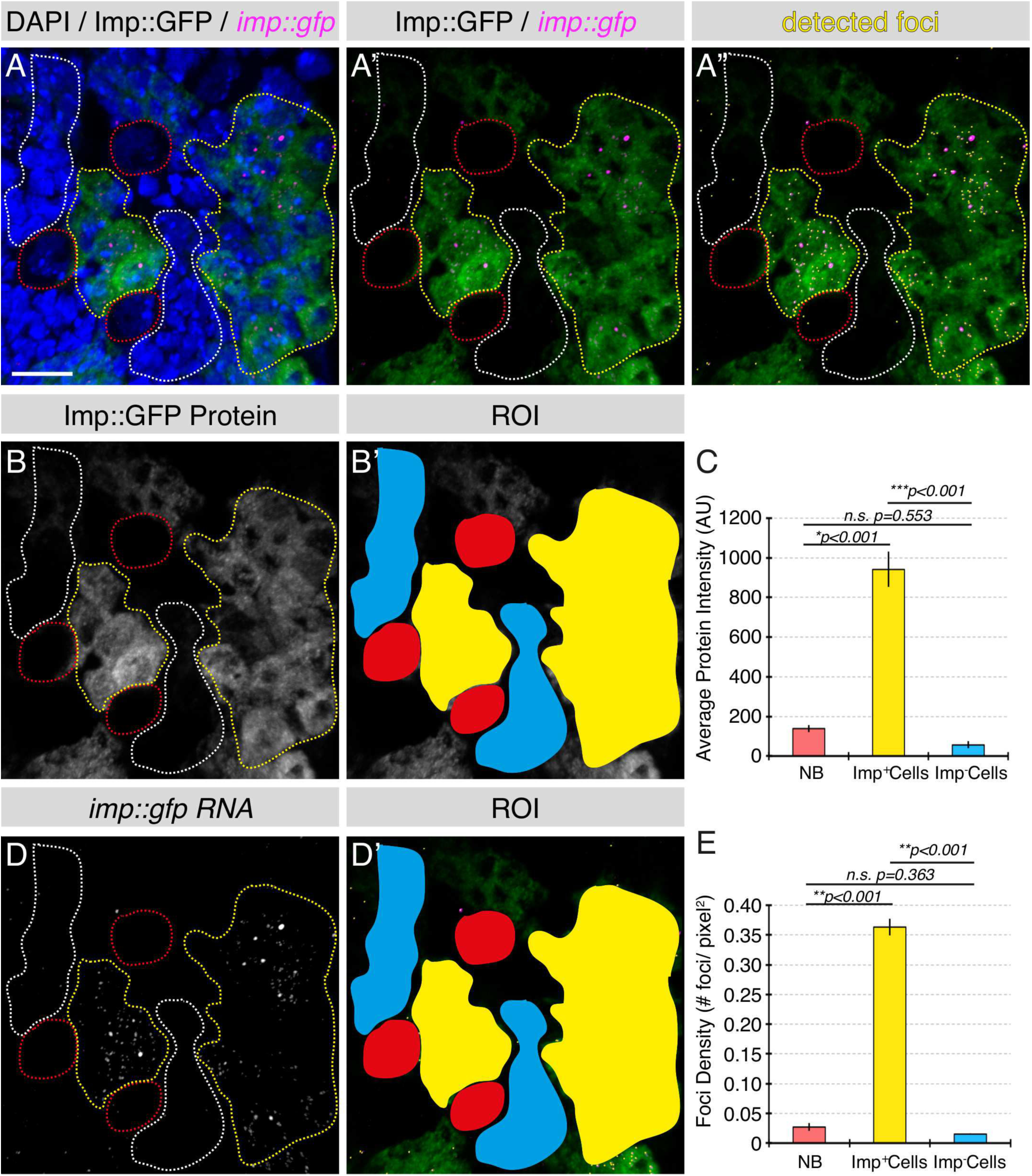
Quantitative analysis of sparse transcripts and RNA/protein expression in neuroblasts and progeny. A-A” *imp* RNA is detected in Imp::GFP larval brains using an smFISH probe targeting the *gfp* sequence (magenta), with simultaneous detection of Imp::GFP protein (green). Individual foci of textitimp mRNA are detected using the “Spots” tool in Imaris (yellow). B-C Imp protein is expressed at a low level in third instar larval neuroblasts and is selectively up-regulated in a sub-population of neuroblast progeny. Quantitative analysis shows Imp protein level is significantly higher in selected neuroblast progeny (C). D-E Up-regulation of *imp* mRNA is found in the cell population that also expresses a high level of Imp protein (D-D’: yellow dotted line region). Quantitative analysis of foci density reveals that neuroblast progeny with higher levels of Imp protein also exhibit a significantly increased level of *imp* mRNA (E). neuroblast: red; Imp+ progeny: yellow Imp-progeny: white. n= 3 cells or cell clusters/brain, 3 brains total. Scale bar represents 10μm.

## 4. Results

Exemplary images of data produced using the protocol above are shown in Figure 2 (RNA dual color detection) and Figure 3 (smFISH combined with antibody staining). To demonstrate single transcript detection in deep brain structures we show smFISH labelling of *msp-300* mRNA in the larval mushroom body calyx [18-19]. The adult *Drosophila* brain is also a heavily researched system with relatively untapped potential for investigating post-transcriptional gene regulation. Therefore, we also include an example of our smFISH protocol in the adult brain targeting *CamKII*, an mRNA known to be compartmentally localized in neurons [20] and whose protein product has an established role in neural plasticity [21].

### 4.1 Application of smFISH to the study of post-transcriptional regulation in the *Drosophila* central nervous system

The proliferative potential of neuroblasts, as well as the specification of the neuroblast progeny fate, requires genes to be expressed at the correct level in the appropriate cell at a specific time during development. Currently, the general focus has been on temporal- and spatial-specific gene regulation at the level of transcription [22-25]. Using the smFISH method described above, we can now rigorously test this hypothesis. With smFISH probes designed against the exon and intron of the gene in question, it is possible to detect mature cytoplasmic mRNA and distinguish it from nuclear nascent transcripts that are detected by intron probes as very bright foci. Such nascent transcript foci consist of primary transcripts decorating the gene locus and quantitating their fluorescence intensity provides a measure of the level of primary transcription. Similarly, quantitating the levels of cytoplasmic signal from exon probes provides a direct measure of the mature transcripts after they are transcribed and exported from the nucleus. Comparing both the level of cytoplasmic mRNA and the level of transcription between cells or in different conditions provides insight into how the gene is regulated, thus providing a quantitative tool to measure mRNA stability and other mechanisms of post-transcriptional regulation.

Here, we use two extensively studied genes, brain tumor (*brat*) and IGF-II mRNA-binding protein (*imp*), as examples. Both Brat and Imp are RNA binding proteins and key regulators of neurogenesis in *Drosophila*. *brat* mutant larvae form supernumerary neuroblasts and brat mutant clones were found to show unregulated cell proliferation [26-28]. Antibody staining, as well as quantitative PCR following cell sorting, have shown that Brat protein and *brat* RNA are both up-regulated in neuroblast progeny relative to neuroblasts, but how this is achieved is not known [14] [29]. Expressing Imp at the correct time during neuroblast development is essential for the specification of neuroblast progeny fate [30], and overexpressing Imp results in dedifferentiation of progenitor cells [31]. However, the mechanism of upstream regulation for achieving correct Imp expression in different types of cells and developmental stages remain to be elucidated. We demonstrate here the use of smFISH to investigate the upstream regulation of both *brat* and *imp*.

### 4.2 Analysis of transcription level in neuroblast and neuroblast progeny

To distinguish whether the up-regulation of Brat in neuroblast progeny is regulated either at the transcriptional or post-transcriptional level, probes targeting the intron and exon region of the gene were used for smFISH experiments (Fig. 4A). In order to analyze the level of transcription, primary transcription foci were detected using the intron probe set (Fig. 4A, B-B’). Images were imported into Imaris image analysis software and transcription foci were automatically identified using the “Spots” tool (Fig. 4B”). Users are required to input an estimated diameter of the foci to be identified; for which we use the width of the point spread function given by the diffraction limit of the dye’s excitation wavelength (λ/2NA, λ=wavelength, NA=numerical aperture) [32]. The intensity of the identified foci was then exported for further analysis. In the current example, the intensity of the brat transcription foci in neuroblasts is compared with that of the neuroblast progeny. Statistical analyses show that there is no significant difference between the levels of brat transcription in neuroblasts compared to their progeny (Fig. 4C).

### 4.3 Analysis of total transcript level using average intensity

To test whether the total *brat* mRNA level is up-regulated in neuroblast progeny, we analyzed the average intensity of *brat* transcripts detected using the brat exon probe set (Fig. 4 A, D-D’). The images were analyzed using the free image analysis software ImageJ. A maximum intensity projection of the acquired image was generated using the “Z Project” tool. Independent projections were then created for neuroblasts and neuroblast progeny clusters in order to accurately capture the 3D space that the cell/cell cluster occupies. From the projection image of the cell (neuroblast) or cell cluster (neuroblast progeny), a user-defined binary mask was drawn for regions of interest (ROIs) and the average intensity of each ROI was measured (Fig. 4D”). In the case of brat, the level of total *brat* mRNA is significantly increased in neuroblast progeny, consistent with previously published qPCR results (Fig. 4E; [4]). Taken together with the analysis of the transcription level analysis presented in section 4.2, it is clear that up-regulation of the brat gene in neuroblast progeny is controlled at the post-transcriptional level, not by transcription.

### 4.4 Analysis of total transcript level by foci counting

An alternative to using average intensity as a measure of total transcript level is to count the number of individual foci, particularly for transcripts with sparse expression. Here we investigate the upstream regulation of Imp expression as an example.

We aimed to address whether the cell type specific Imp protein expression is regulated by its mRNA level. If Imp protein and mRNA level correlate with each other, this would suggest Imp expression is regulated at the pre-translational level. Alternatively, if Imp protein and mRNA level do not correspond, this would suggest translation or post-translational regulation. We first quantified the level of Imp protein in neuroblasts and their progeny with either high or low Imp expression (Fig. 5A-C). Statistical analyses show Imp protein expression is low in neuroblasts and is expressed only in a subpopulation of neuroblast progeny (Fig. 5C). Next, we analyzed the level of *imp* mRNA in these three cell groups using smFISH probes targeting the GFP open reading frame. As the expression level of the imp transcript is sparse, the “Spots” tool in Imaris could be used to detect each of the individual foci. Subsequently, the number of foci in each ROI was counted and the total level of transcript expression was quantified by calculating foci density for each selected cell population (foci density = number of foci/area of ROI) (Fig. 5D-E). We found the pattern of *imp* expression level closely mirrored that of Imp protein. From these data, we conclude that unlike *brat*, the cell-type-specific Imp expression level is regulated at the level of transcription.

### 4.5 Using smFISH to identify neuroblast, ganglion mother cells, and immature neurons

In order to study neural development, reliable labeling of the different cell types in the brain is essential. This is most commonly accomplished with antibody staining. However, immunohistochemistry (IHC) requires high-quality antibodies that provide adequate signal-to-noise on fixed tissues. For *Drosophila*, high-quality antibodies are rarely available commercially and are not easy to produce. Choice of antibody combinations is also limited by cross-species reactivity. In this section, we present the use of smFISH as a simple and time-efficient alternative.

The most frequently used neuroblast label in the larval CNS is Deadpan (Dpn) and the label for young neuroblast progeny, also known as ganglion mother cells (GMCs), is Asense (Ase) (Fig. 6A-A”’). Since Ase protein is also expressed in the neuroblasts, it is best that Ase is used in conjunction with Dpn for GMC labeling (GMC: Ase+ Dpn-) (Fig. 6A’,A”’). To overcome the problem of sourcing suitable antibodies, we have developed an alternative labeling regime using smFISH probes targeting *cyclin B*, which labels neuroblasts (Fig. 6C-C’), and *castor*, which labels GMCs (Fig. 6D-D’). By choosing suitable fluorescent labels, these probes can be used together or independently (and are compatible with antibody labelling for additional markers).

**Figure 6.**
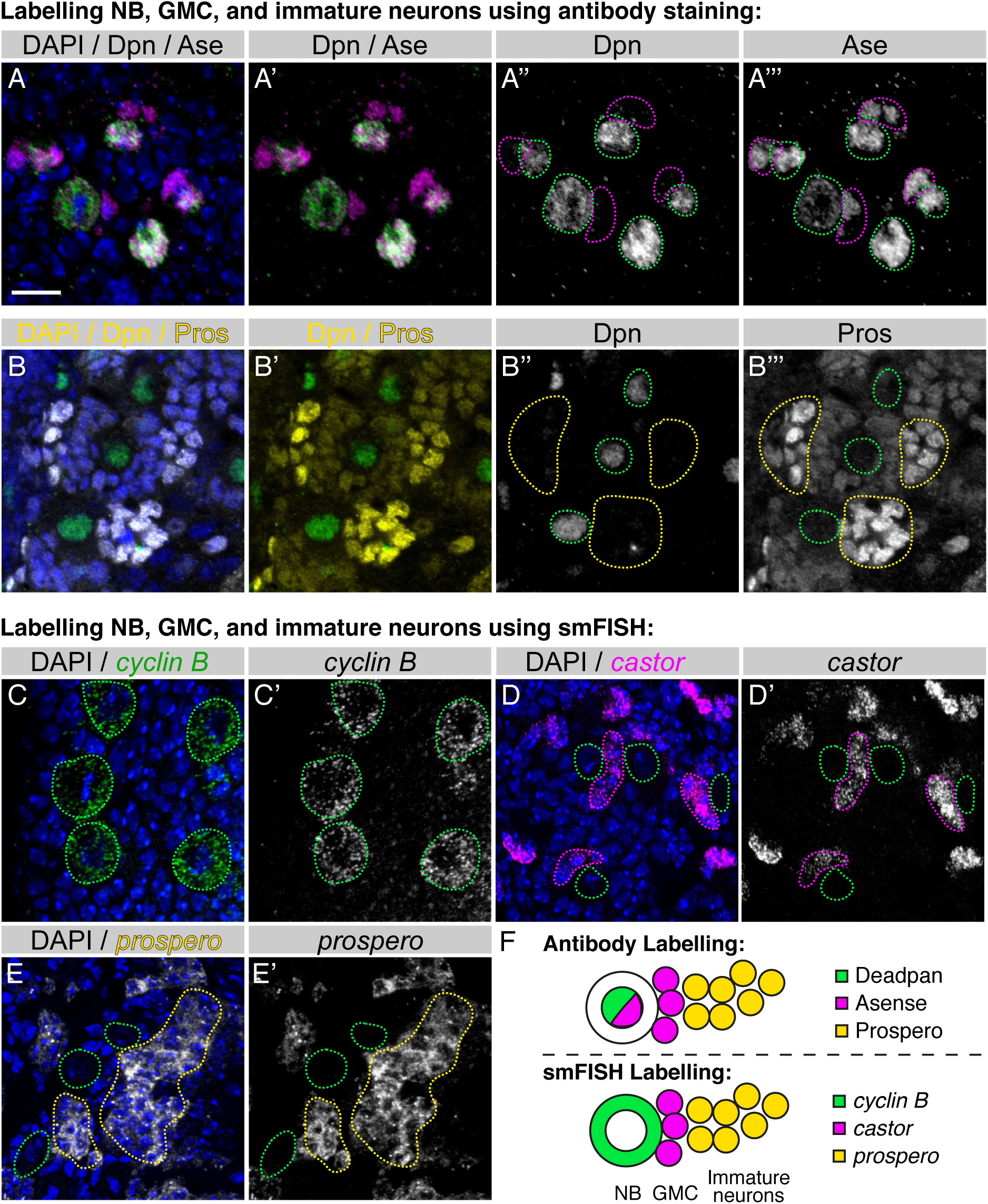
Using smFISH to identify neuroblasts, GMCs and immature neurons. A-A”’ Conventional antibody labelling for neuroblast (Deadpan antibody; green) and ganglion mother cells (GMCs) (Deadpan-Asense+; magenta). B-B”’ Immature neurons are distinguished from the newly born neuroblast progeny by the up-regulation of Prospero protein (bright yellow cells: B-B’; yellow outlined region: B"-B”’). C-E’ Labelling neuroblasts, GMCs and immature neurons using smFISH. Neuroblasts, GMCs and immature neurons are labelled with *cyclin B*, *castor* and *prospero*, respectively. F Schematics showing labelling of neuroblast and neuroblast progeny using the conventional antibody staining method or the new smFISH method. neuroblast: green outlined region; GMCs: magenta outlined region; immature neuron: yellow outlined region. Scale bar represents 10μm.

Immunofluorescence labeling of immature neurons in the third instar larval brain has also been challenging. Antibodies against the commonly used immature neuron label, Elav, tend to have poor signal-to-noise, and Prospero (Pros) protein is expressed in both GMCs (low levels) as well as immature neurons (high levels; Fig. 6B-B”’). We have found that a smFISH probe set that targets the 3’UTR region of the *pros* transcript is an effective method to label immature neurons (Fig. 6E-E’). This label generates a minimum signal in neuroblasts and GMCs with high signal in immature neurons. As with other smFISH labels described in this manuscript, the pros probe set is compatible with conventional IF labelling. Collectively, we propose the use of smFISH with probes targeting *cyclin B*, *cas* and *pros* as a new and more effective method for neu-roblast, GMC and immature neuron labelling compared to the traditional Dpn, Ase and Pros antibody staining method (Fig. 6F).

## 5. Concluding remarks

In summary, our modified smFISH protocol offers a range of tools for studying post-transcriptional gene regulation in complex intact tissues. The resulting images have a high signal to background ratio even when imaging at a depth of 40μm and have the sensitivity to detect rare single transcripts, i.e., fewer than 100 transcripts per cell. We demonstrate the use of our technique to quantitate the brightness of nascent transcription foci and cytoplasmic mRNA levels at sub-cellular resolution. Such data provides a way to investigate post-transcriptional mechanisms at the single cell level within intact complex tissues. Our smFISH protocol is rapid and straightforward, with a small number of reagents and steps, while remaining adaptable for use with antibody staining and the simultaneous detection of multiple RNAs. Our optimized protocol on whole *Drosophila* brains demonstrates the application of sm-FISH as a tool for post-transcriptional regulation and RNA biology in thick tissue. We also show that smFISH can be effectively used to mark specific cell types in addition to, or as a replacement for, cell specific antibody labelling.

## Acknowledgements

We are grateful to Alan Wainman and Richard Parton for advice on advanced microscopy. We would also like to thank the Bloomington Drosophila Stock Center and Jurgen Knoblich for fly stocks and antibodies. L.Y. was funded by the Clarendon Trust and Goodger Fund. D.I.H. was funded by University College London. S.W. is funded by a Wellcome Trust Principal Research Fellowship in the Basic Biomedical Sciences and the Bettencourt-Schueller Foundation. J.T., D.E., and I.D. were funded by a Wellcome Senior Research Fellowship to I.D. (081858), MICRON Oxford http://micronoxford.com, supported by a Wellcome Strategic Awards to ID (091911/B/10/Z and 107457/Z/15/Z).

## Notes

1 Care should be taken to ensure that formamide is not allowed to oxidise. Un-opened bottles should be stored at 4°C. Once opened, the liquid should be immediately dispensed into 1 ml aliquots in a fume hood, flash frozen in liquid nitrogen, and stored at -70°C.

2 Fresh SSC solution is essential for achieving a good signal-to-noise ratio of the smFISH experiments. 20x SSC solution can be stored at room temperature for several weeks.

3 In comparison to other published FISH protocols, we find no additional benefit in using RNAse inhibitors or non-specific blockers, e.g., salmon sperm DNA or tRNA.

4 Aliquots of blocking buffer should be stored at -20C. Care should be taken to ensure blocking buffer is prepared in a sterile environment (i.e. under laminar flow hood).

5 Washes should be carried out in 0.3% PBST instead of PBSTX, as Triton-X can affect tissue morphology.

6 We typically dilute the probe concentration to 1μM (1:50 dilution from stock solution). For genes with low expression level, the probe may need to be used at a concentration of 2-5μM. We recommend testing a range of probe concentrations (between 0.1-5μM) for genes being detected for the first time.

7 Hybridisation step should not be longer than 15 hours as long incubations greatly reduce the signal-to-noise ratio.

8 Care should be taken to avoid pipetting brains into pipette tip as the brains have a high tendency to adhere to the inside wall of the tip after the hybridisation step.

9 For Type I neuroblasts, mount brains with ventral side facing down towards the coverslip. For Type II and MB neuroblasts, and for MB neuropil in the larva or adult brain, mount brains dorsal side down to ensure the cell type of interest is closest to the coverslip (Fig. 1C).

10 Place two pieces of double-sided tape approximately 20mm apart. This will secure the coverslip in position.

